# Mutations in *iclR* increase evolvability by facilitating compensation that exposes cryptic beneficial mutations in experimental populations of *Escherichia coli*

**DOI:** 10.1101/2024.02.21.581449

**Authors:** Rachel K Staples, Tim F. Cooper

## Abstract

Evolvability describes the potential of a population to generate beneficial variation. Several mechanisms that increase evolvability have been demonstrated, including the action of systems that reveal accumulated beneficial variants following an environmental shift. We examine the basis of an increase in the evolvability of *Escherichia coli* lines that were first selected in an environment supplemented with glucose as sole carbon source and then transferred to an otherwise identical lactose supplemented environment. These lines increased in fitness significantly more quickly in the lactose environment, and reached a higher final fitness, than did naïve ancestral lines. In four of six lines this increased evolvability can be explained by mutations in *iclR* that were selected in glucose but were significantly deleterious in lactose, masking the effect of other generally beneficial mutations. Secondary mutations that compensated for this cost resulted in large fitness increases. We did not detect any consistent genetic signature associated with the compensation, suggesting that different pathways were responsible and, therefore, that it can occur at a relatively high rate. That mutations selected in one environment will become deleterious following an environmental shift, so that compensation provides potential for a large subsequent fitness increase represents a potentially common and general mechanism of evolvability in changing environments.

## Introduction

Evolvability describes the ability of a biological system to produce heritable adaptive variation (1). This ability can vary between evenly closely related genotypes and can readily evolve to impact evolutionary outcomes (2–11). Highly evolvable genotypes can be thought of as being in regions of a fitness landscape from which increased fitness can occur through many pathways or to high peaks (1, 12, 13). This situation can occur through general underlying mechanisms, for example effecting the generation or maintenance of population genetic variation through changes in mutation rates or mutation effects (1, 14–16). Evolvability can also depend on specific interactions, for example the evolution of a citrate utilization phenotype that depends on specific potentiating mutations (17–19).

The best studied mechanism of increased evolvability is an increase in the genomic mutation rate that can provide an advantage by generating a larger number of new mutations for selection to act upon, increasing the potential for adaptive evolution (20–23). However, the benefits of a mutator phenotype are dependent on context and environment. While high mutation rates can increase genetic variation, genetic diversity, and potential for adaptation, they can also increase the risk of deleterious mutations and genetic instability (24). Therefore, the balance between the benefits and costs of high mutation rates may be specific to genotypes and their selective environment. For example, whereas fluctuating environments provide a backdrop of consistent fitness improvement providing opportunity for mutator selection, linkage with deleterious mutations may confer long-term costs (24).

Other proposed mechanisms of evolvability involve changes that allow the expression of cryptic preexisting genetic variation (5, 16, 25–27). Such mechanisms allow for the accumulation of neutral genetic variation until that variation is revealed, often by exposure to a stressful environment, with the chance of including some new adaptive response to the environmental challenge. For example, in times of environmental stress, the Hsp90 protein chaperone may become overwhelmed and fail to properly stabilize proteins, releasing accumulated genetic variation to be expressed and for selection to act upon (28).

A similar mechanism that involves compensation of conditionally deleterious mutations was proposed in Phillips et al. (2016). In principle, mutations selected in one environment could remain beneficial, or become neutral or deleterious following an environmental change (29, 30). Considering the combined effect of multiple mutations that are substituted prior to an environmental change, if some of them are deleterious in a new environment their overall effect on fitness will be less than would otherwise be the case. Indeed, it is possible that mutations that are deleterious in a new environment could counteract the positive effects of beneficial alleles, masking the underlying potential of an otherwise more fit genome. Compensatory mutations that specifically neutralize the negative effects of conditionally deleterious alleles, exposing the positive fitness effects of the pre-existing beneficial alleles, could provide a mechanism to rapidly increasing fitness, thereby increasing evolvability (3, 31).

We previously reported an experiment in which six replicate populations founded from a single *E. coli* ancestor were selected for 2,000 generations in a glucose-limited environment (3, 32). A single clone from each of the glucose-evolved populations, and the ancestor, were used to found new replicate populations that were evolved in lactose for 1,000 generations (Fig. 1) (3). Although the glucose-evolved founders were initially no more fit than their ancestor in the lactose environment, they were found to be more evolvable because their fitness consistently increased more rapidly, and to a higher level, than did populations evolved from their ancestor (32). The ancestor and glucose-evolved clones had the same mutation rate, suggesting that the genetic basis of greater evolvability in lactose was a specific consequence of glucose adaptation.

**Figure 1.**
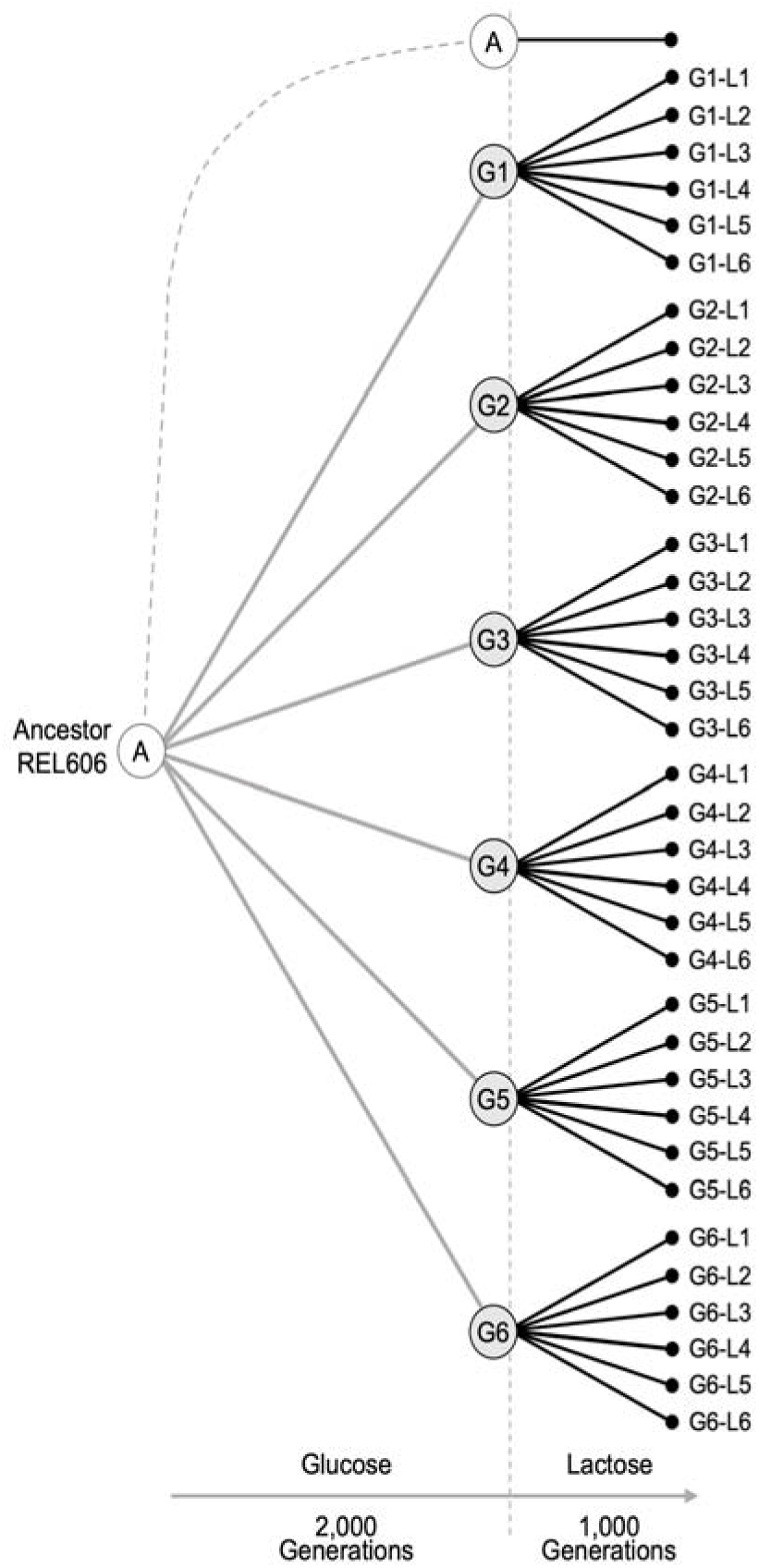
Schematic of experimental evolution design. A single ancestor strain (‘A’) was previously used to found six replicate populations that evolved in a glucose environment for 2,000 generations. After that, a single clone was isolated from each of the glucose evolved populations (G1-6). These six glucose-evolved founders were each used to found six replicate populations that were evolved for 1,000 generations in a lactose environment. After lactose evolution, a single clone was isolated from each evolved population for sequencing. A single lactose evolved clone from each founder was used for additional genetic manipulation and fitness estimates. Lactose evolved clones are denoted by their glucose-evolved founder and the replicate lactose population they were isolated from.

A candidate basis for prior glucose selection increasing evolvability in lactose are mutations in *iclR,* encoding the isocitrate lyase transcriptional regulator that controls expression of enzymes involved in the glyoxylate cycle. Different *iclR* mutations were found in four of the six glucose-adapted founders but not in any of the populations evolved from the same ancestor and selected only in environments containing lactose as the sole carbon source (33). This pattern suggests the substituted mutations in *iclR* are beneficial in a glucose-limited environment but are neutral or deleterious in a lactose-limited environment. If they are deleterious in lactose, compensation for that effect, could serve to reveal generally beneficial mutations that accumulated during glucose selection, providing a mechanism for more rapid adaptation and greater evolvability. We test this hypothesis by determining the fitness effect of evolved *iclR* alleles in the clones used to found the lactose selected populations and in evolved clones isolated from the end of the experiment. Consistent with our hypothesis, *iclR* mutations tended to be deleterious at the start of the experiment but were neutral by the end, indicating that deleterious effects had been compensated thereby unmasking the beneficial effect of other mutations that accumulated during glucose adaptation.

## Materials and methods

### Bacterial strains

Populations evolved for 2,000 generations in an environment consisting of Davis-Mingioli (DM) minimal medium supplemented with glucose as the sole carbon source has previously been described (3). Briefly, a single ancestor strain, *E. coli* REL606, and an otherwise isogenic derivative able to utilize arabinose, REL607, were used to create six replicate populations that were evolved under glucose selection for 2,000 generations (32). A single clone was isolated from each of the six populations (Table S1). Each of these six glucose-adapted clones (denoted G1-6, where the number denotes the source population) were used to found six new replicate populations that were selected for 1,000 generations in the same minimal medium environment used for the initial period of evolution except supplemented with lactose as the sole carbon, yielding a total of 36 ‘G-L’ populations (Fig. 1). A single clone was isolated from each G-L evolved population and stored at −80°C for subsequent analysis and sequencing. G-L populations are referred to using the general nomenclature GX-LY with ‘X’ indicating the founder glucose selected clone and ‘Y’ indicating the replicate lactose-selected population started from that founder.

### Strain construction and replacement of *iclR* alleles

Founder clones G2, G3, G4, and G6 were previously identified as having mutations in *iclR* (3). We added these evolved *iclR* alleles into the ancestral strain and reverted evolved *iclR* alleles in G2, G3, G4, and G6, and in clones isolated from evolved populations started from each of these founders. To do this we constructed allele exchange vectors by adding a DNA region containing a relevant part of the *iclR* gene into the suicide-plasmid pDS132 (34) (Tables S2 and S3). Constructed plasmids were used to transform MFD*pir* cells with selection on solid lysogeny broth (LB) medium supplemented with 25 µg/ml chloramphenicol (Cm) and 0.3 mM diaminopimelic acid (DAP; which MFD*pir* requires for growth) (35). Presence of the cloned insert was confirmed by PCR. Biparental matings were used to transfer *iclR*-containing plasmids into target recipient strains using a previously described protocol (36).

### Fitness competitions and statistical analyses

To measure the fitness effects of the four evolved *iclR* mutations, the common ancestral strain was separately competed with the four founders, the four 1,000 generation G-L clones evolved from those founders, and corresponding constructed derivatives with ancestral *iclR* alleles. A derivative of the ancestor carrying an opposite neutral arabinose marker to that present in all other strains allowed for enumeration of competitors on tetrazolium-arabinose (TA) indicator agar (37).

Replicate fitness competitions were carried out in deep 96-well polypropylene blocks incubated at 37°C with shaking at 200 RPM. All competition strains were inoculated from freezer stocks to 3 mL LB medium and grown overnight at 37°C. Competitors were acclimated to the competition environment by diluting 1:1000 from LB cultures into 1 ml DM medium containing 210 mg/L lactose (DM210 lactose), grown for 24 hours, then diluted 1:100 into fresh DM210 lactose and again grown for 24 hours. Pairwise competitions were performed by combining equal volumes of both competitors, then diluting 1:100 into 1 ml DM210 lactose and incubating for 24 hours. Competition cultures were sampled immediately after mixing and again after 24 hours by diluting and plating onto TA agar. For each replicate competition, the number of colonies representing each competitor were counted and used to calculate relative fitness, *w*, as the ratio of each competitor’s Malthusian parameter (37).

### Whole genome re-sequencing and mutation identification

Thirty-six G-L 1,000 generation evolved clones were sequenced to identify new mutations acquired during lactose evolution. Genomic DNA was isolated and purified using the Wizard Genomic DNA Purification Kit protocol for Gram negative bacteria at one-third volume (Promega), then quantified using SYBR Green I Nucleic Acid Stain (Invitrogen) in a SpectraMax M5 Fluorescence Microplate Reader (Molecular Devices). Libraries were individually created for each clone following a modified Nextera XT DNA Library Prep Kit protocol (Illumina). Tagmentation and amplification was carried out at one-quarter volume with Nextera XT Index Kit v2 adapters, then SPRIselect beads (Beckman Coulter) were used at full volume for fragment size selection. Libraries were individually quantified using the Qubit dsDNA High Sensitivity Assay with a Qubit 2.0 Flourometer (ThermoFisher) and fragment size was confirmed for a representative subset of samples using the Tapestation 4200 (Agilent), after which all libraries were diluted and pooled at a concentration of 1 ng/ul in preparation for sequencing. The pooled sample was sequenced on an Illumina Hiseq X producing 150 base-pair, paired-end reads at the University of Houston Seq-N-Edit Core. Mutations acquired by sequenced clones during lactose selection were identified using the computational pipeline breseq to compare with the previously sequenced ancestral strain and glucose-evolved founders (38).

### Mutational parallelism

Newly identified mutations in 1000 generation G-L clones were screened for an effect of evolutionary history on genetic parallelism (39, 40). Briefly, a parallelism index (PI) was estimated as the mean number of in-common mutated genes identified among all pairs of clones derived from a given founder. Similarly, a mutation convergence index (CI) was estimated as the mean number of mutated genes shared by pairs of clones from different founders. A positive PI:CI ratio indicates relatively higher mutation parallelism among clones of a focal founder, reflecting an impact of founder genotype on the genes mutated during lactose evolution. Mutations occurring in founder clones during selection in glucose were excluded this analysis.

### Statistical analysis

R version 4.3.1 was used for all analysis and plotting (41). Shapiro-Wilk tests were used to confirm whether the relative fitness estimates were normally distributed, and Levene’s tests for homogeneity of variance were performed to determine whether variance across compared groups was equal. Welch’s t-tests were used to compare the relative fitness of strains differing at the *iclR* locus as indicated in the main text.

## Results

### Relative fitness estimates in lactose and glucose

We previously compared rates of adaptation to a lactose environment of replicate lines started from a naïve ancestral strain and six independently derived ‘founder’ clones (3). Founder clones were started from the ancestor and selected in a glucose environment for 2,000 generations before being moved to the lactose environment. The founder clones were initially less fit than the ancestor in the lactose environment but were more evolvable. During 1,000 generations of evolution in the lactose environment, founder derived lines increased in fitness more quickly and to a higher level than did lines started from the ancestor (3).

Sequencing of the six founder clones identified several mutations that were also seen in lactose selected lines and that we presume increase fitness in lactose. We also found mutations in *iclR* in four founders (G2, 3, 5, and 6; referred to as *iclR^Ev^* founders) (3). Mutations in *iclR* were not seen in any population evolved only in lactose (33). We hypothesized that *iclR* mutations were costly in lactose and acted to mask the effect of mutations that were beneficial in both glucose and lactose, and that compensation of this cost during lactose selection increased evolvability by unmasking the effect of the beneficial mutations.

To test this hypothesis, we estimate the fitness effect of evolved *iclR* alleles by replacing them with the ancestral allele and comparing the fitness of initial and constructed strains. These estimates allow us to test the prediction that evolved *iclR* alleles were costly in founder strains but changed to become neutral or beneficial following 1,000 generations of evolution in lactose. We find that evolved *iclR* alleles impose a fitness cost in the lactose environment in all *iclR^Ev^* founders (Fig. 3). This cost was significant in G2, G3 and G6 founders, and marginally non-significant in G5 (Welch’s one-tailed *t*-test: G2: *t_21.6_* = 3.6, P < 0.01; G3: *t*_12.9_ = 9.3, P < 0.01; G5: *t*_12.4_ = 1.7, P = 0.06; G6: *t*_10.9_ = 3.8, P < 0.01) (Fig 3, Tables 1 & S4). Furthermore, founders that were reverted to encode the ancestral *iclR* allele had significantly higher fitness in lactose than did the ancestor, indicating that the net effect of the remaining glucose substituted mutations was positive in lactose (Fig. 3, Table 1 & S4).

**Figure 2.**
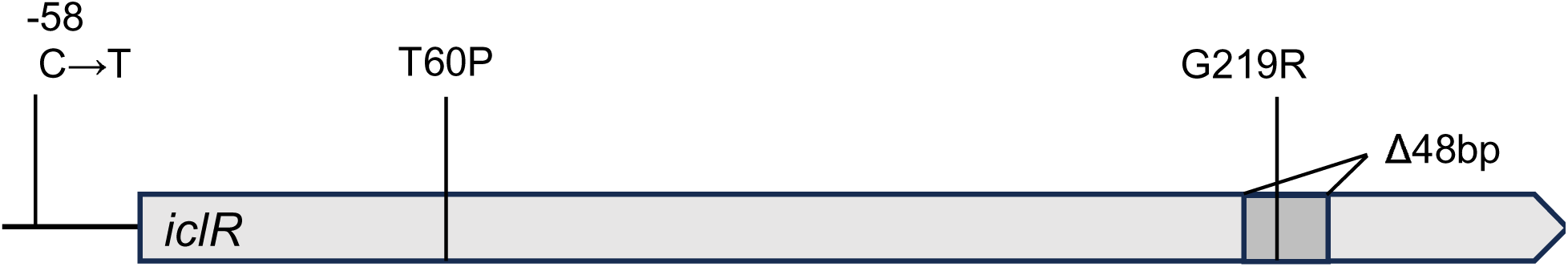
Schematic of *iclR* gene with approximate locations of the four mutations identified in glucose-evolved founders G2, G3, G5, and G6.

**Figure 3.**
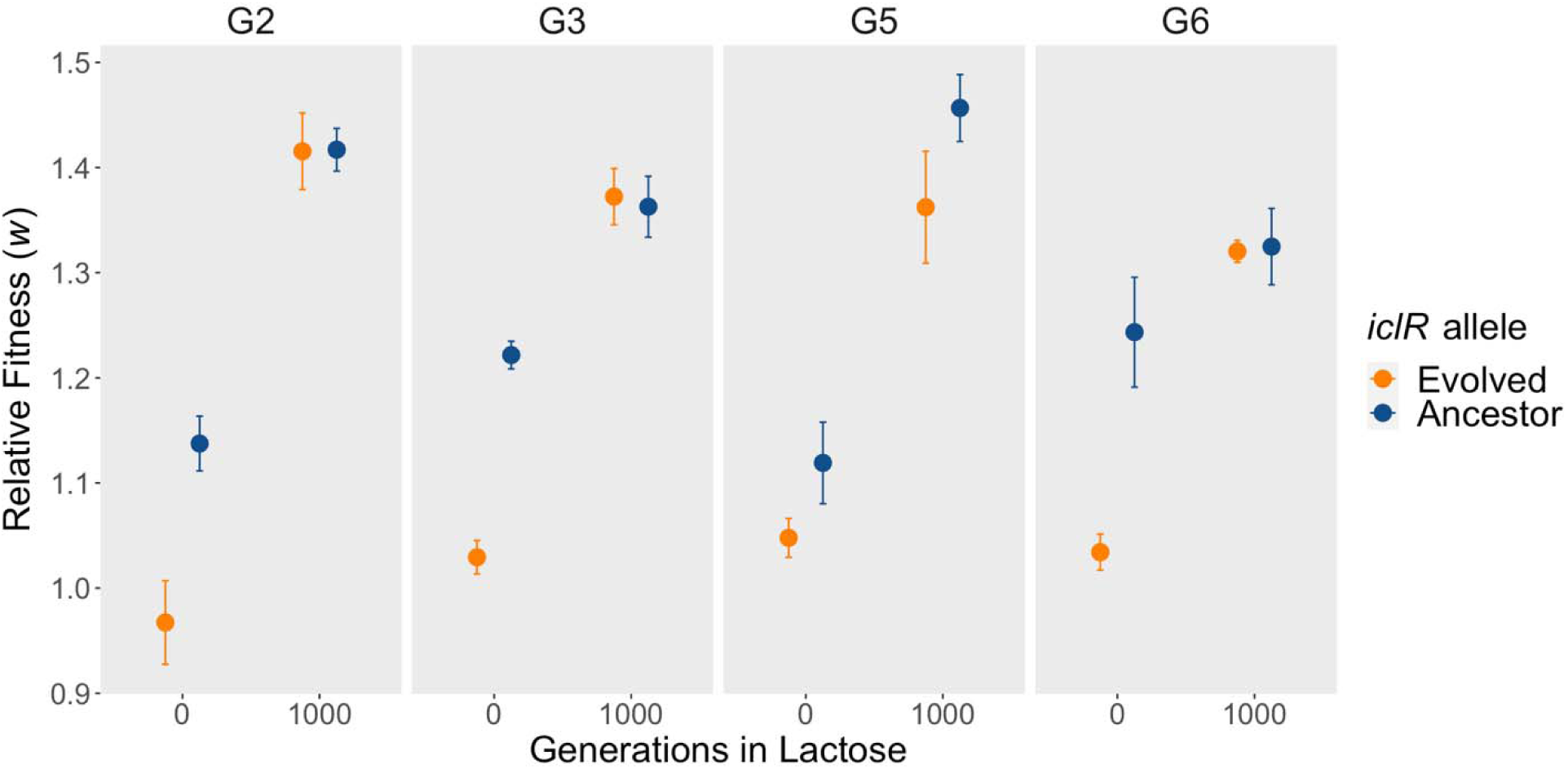
Fitness measured in lactose of strains with and without evolved *iclR* alleles in glucose-evolved founders and one lactose evolved clone derived from each founder. Symbols indicate the mean of five or more independent replicates. Error bars represent standard error of the mean. Relative fitness values equivalent to 1.0 indicate a strain is equally as fit as the ancestor, values greater than and less than 1.0 indicate more, or less fit than the ancestor, respectively.

**Table 1.** Fitness of founder, evolved and constructed strains in lactose and glucose.

|  | Strain | Gen. in<br>lactose | *iclR*<br>allele | Mean Relative Fitness (*w*) |  |  |  |
| --- | --- | --- | --- | --- | --- | --- | --- |
|  |  |  |  | Lactose | 95% CI | Glucose | 95% CI |
| Founder | G2 | 0 | T60P | 0.967 | 0.172 | 1.201 | 0.105 |
|  | G2-*iclR*<sub>anc</sub> | 0 | Anc | 1.137 | 0.123 | 1.250 | 0.112 |
|  | G2-L1 | 1,000 | T60P | 1.415 | 0.178 | – | – |
|  | G2-L1-*iclR*<sub>anc</sub> | 1,000 | Anc | 1.417 | 0.096 | – | – |
| Founder | G3 | 0 | G219R | 1.029 | 0.075 | 1.186 | 0.098 |
|  | G3-*iclR*<sub>anc</sub> | 0 | Anc | 1.222 | 0.073 | 1.150 | 0.124 |
|  | G3-L1 | 1,000 | G219R | 1.372 | 0.123 | – | – |
|  | G3-L1-*iclR*<sub>anc</sub> | 1,000 | Anc | 1.363 | 0.134 | – | – |
| Founder | G5 | 0 | -58/-142 | 1.048 | 0.103 | 1.208 | 0.085 |
|  | G5-*iclR*<sub>anc</sub> | 0 | Anc | 1.119 | 0.179 | – | – |
|  | G5-L1 | 1,000 | -58/-142 | 1.362 | 0.295 | – | – |
|  | G5-L1-*iclR*<sub>anc</sub> | 1,000 | Anc | 1.457 | 0.177 | – | – |
| Founder | G6 | 0 | Δ48 | 1.034 | 0.078 | 1.210 | 0.145 |
|  | G6-*iclR*<sub>anc</sub> | 0 | Anc | 1.243 | 0.241 | 1.248 | 0.260 |
|  | G6-L1 | 1,000 | Δ48 | 1.320 | 0.066 | – | – |
|  | G6-L1-*iclR*<sub>anc</sub> | 1,000 | Anc | 1.325 | 0.202 | – | – |
| Ancestor | Anc-*iclR*<sub>G2</sub> | 0 | T60P | 0.981 | 0.035 | 1.000 | 0.057 |
|  | Anc-*iclR*<sub>G3</sub> | 0 | G219R | 1.002 | 0.021 | 0.999 | 0.098 |
|  | Anc-*iclR*<sub>G5</sub> | 0 | -58/-142 | 0.984 | 0.019 | 1.023 | 0.067 |
|  | Anc-*iclR*<sub>G6</sub> | 0 | Δ48 | 0.966 | 0.031 | 1.004 | 0.111 |

To test the second part of our hypothesis, that the greater evolvability of the glucose evolved *iclR^Ev^* founder clones was linked to compensation of the cost of evolved *iclR* alleles unmasking the effect of generally beneficial mutations, we reverted evolved *iclR* alleles in four clones isolated from populations started from different *iclR^Ev^* founders and that were evolved in lactose for 1,000 generations. Evolved clones were isolated from lactose evolved lines G2-L1, G3-L1, G5-L1 and G6-L1. Consistent with our prediction, in these evolved clones the evolved *iclR* alleles no longer conferred any significant cost to lactose fitness (Fig. 3, Tables 4 and S4; Welch’s one-tailed *t*-test: G2-L1: *t_11.1_*= 0.04, P = 0.49; G3-L1: *t*_17.9_ = 0.24, P = 0.40; G5-L1: *t*_8.2_ = 1.5, P = 0.08; G6-L1: *t*_5.8_ = 0.12, P = 0.45). Together these results indicate that the evolved *iclR* alleles were costly at the start of the lactose-evolution and that, at least in the surveyed lines, these costs were compensated for during the 1000 generations of lactose evolution.

For completeness, we also added each of the four evolved *iclR* alleles directly to the ancestral background to examine their individual fitness effect in glucose and lactose. We found that none of the evolved alleles had a significant impact on fitness in either environment (Table 1). We expected that the evolved alleles would be beneficial in glucose because they were identified in four of the six glucose-evolved populations, so this result suggests that the influence of *iclR* alleles may be conditional on other mutations arising during the period of glucose selection.

### Whole genome sequencing results

To identify mutations that could be compensating for the deleterious effects of evolved *iclR* alleles, we sequenced evolved clones isolated from each of the six independent lines started by each of the six founders. By comparing genome sequences of these evolved clones to those of the corresponding founder we could determine mutations that arose during evolution in lactose (Fig. 4). Two of the sequenced clones did not share mutations with their respective founders and were excluded from further analysis.

**Figure 4.**
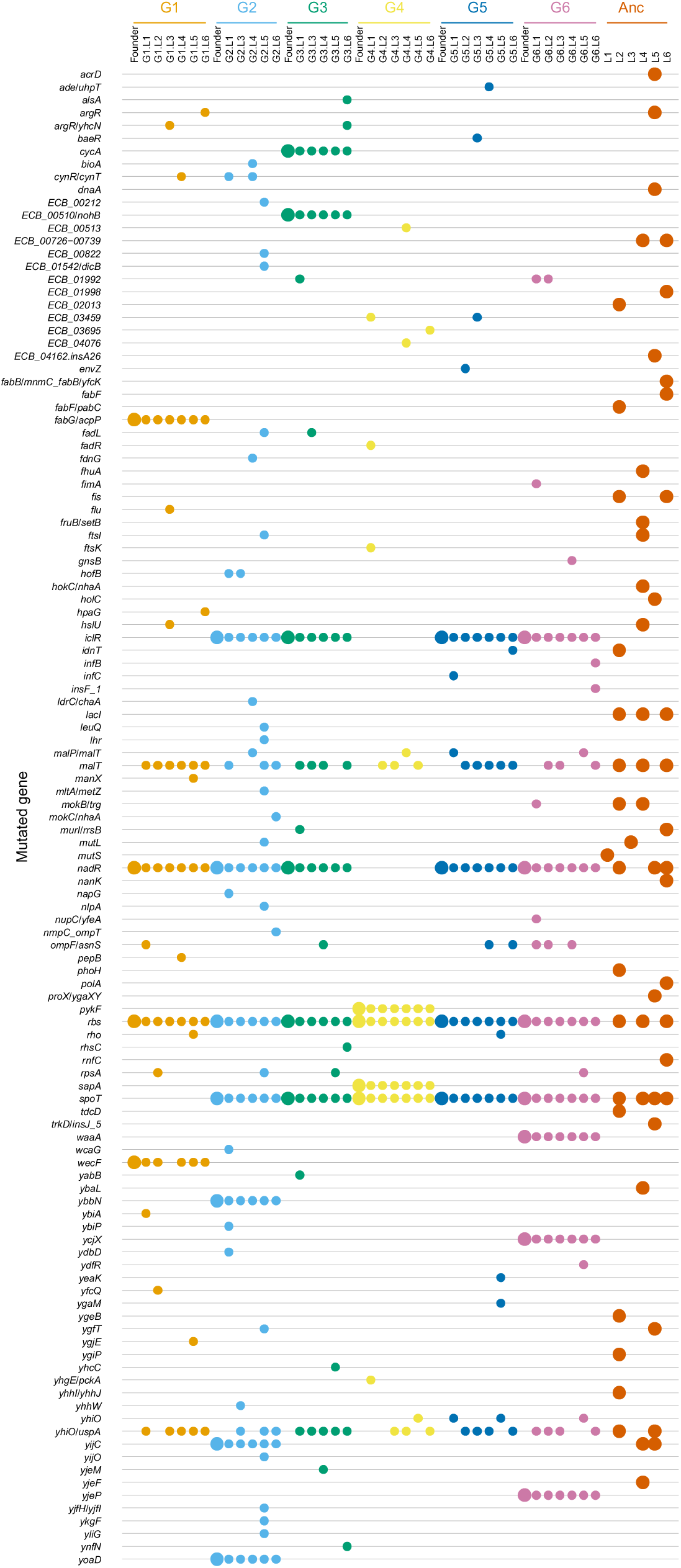
Mutations identified in glucose-evolved founders (G1-6) and the G-L clones evolved from them, after 1,000 generations of lactose evolution. Color of symbols follow the color of the progenitor founder strain, as indicated at the top of the figure. Large symbols indicate founder strains, small symbols indicate derived clones isolated following evolution in lactose.

The number of new mutations occurring during lactose evolution varied among G-L clones, with a median of four new mutations. Most clones had acquired somewhere between one and seven new mutations, although one clone, G2-L5, had 18 new mutations. Among them was a +3 bp insertion in the gene *mutL.* In *E. coli*, the MutL protein is involved in mismatch repair and a mutation within its coding sequence may explain the high number of mutations in this clone (42).

Notably, none of the sequenced clones were found to have acquired any new mutations in the *iclR* gene during lactose evolution, indicating that the change in effect of evolved *iclR* alleles was due to the action of newly arising mutations in genes that directly or indirectly interact with *iclR* (Fig. 5). No new *iclR* mutations were found in clones isolated from the G1 and G4 glucose-evolved populations, consistent with mutations in this gene not being beneficial in the lactose selective environment (Fig. 5). However, a single nonsynonymous mutation in *fadR* was identified in clone G4-L1, which evolved from a founder without any *iclR* mutations. FadR, which regulates fatty acid biosynthesis and degradation, has also been recognized as a proximal activator of *iclR* (Gui et al., 1996). Also of note, mutations in *lacI*, the *lac* operon negative regulator, have commonly been identified in populations selected in lactose environments but no *lacI* mutations were found among these G-L clones after 1,000 generations in lactose (33, 43).

**Figure 5.**
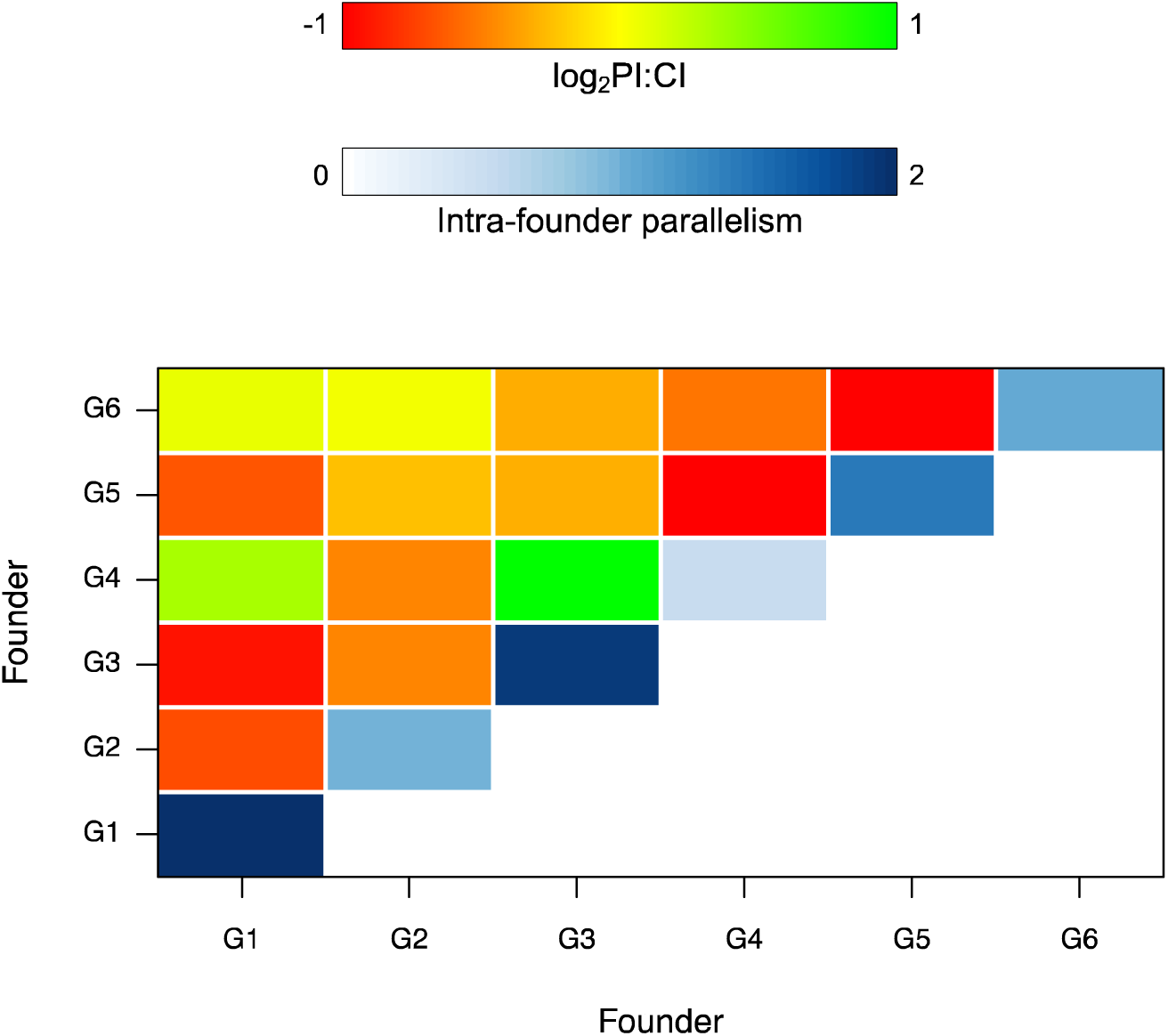
Overlap in mutated genes occurring in different founders. The mean number of shared mutated genes between replicate lines started from each founder (i.e., parallelism) is represented along the blue diagonal. The PI:CI metric is estimated as the ratio of the mean number of shared mutated genes among clones derived from each pair of compared founders over the mean number of shared mutated genes estimated across all clones derived from those founders.

### Mutational parallelism and convergence

To identify mutations that might compensate for the cost of evolved *iclR* mutations, we tested for gene-level mutational parallelism. Parallel mutations that are present in lines started from the founder clones and absent from lines started from the naïve ancestor are candidates for increasing evolvability. Similarly, parallel mutations specific to lines started from the four *iclR^Ev^* founders are candidates for interacting with that mutation to reduce its cost.

Six different genes (including flanking intergenic regions) were found to have mutations in clones isolated from more than one *iclR^Ev^*founder. Multiple distinct mutations were identified in the intergenic *cynR/cynT* and *ompF/asnS* regions, within the coding sequence of *ECB_01992* and *yhiO*, and within coding and intergenic regions of both *argR* and *malT*.

Of note, 28 mutations were identified in the transcription regulator *malT* and its upstream region. These mutations occurred in lines started by all six founders, representing over 75% of all sequenced evolved clones. Four different mutations were identified in *malT* associated intergenic regions and 24 were found in the *malT* coding sequence, of which 18 were unique. Similarly, mutations were identified in the coding or upstream region of the gene *yhiO* in 28 clones representing all six founders. In this case, the majority of mutations (24/28) were 2-3 base pair (bp) insertions located in the intergenic region, but 2 different +3bp insertions and 2 nonsynonymous substitutions were also identified within the coding sequence. This broad gene-level parallelism, and the fact that *malT* and *yhiO* mutations were not identified in the glucose-evolved founders, strongly suggests that they are specifically beneficial in lactose. However, mutations in both genes occur in multiple lines started from *iclR^Ev^* and *iclR^Anc^* founders, and from lines started directly from the ancestor. This suggests both that they are not responsible for compensation of the *iclR* mutations and that they do not explain greater evolvability of the glucose founder started lines (Fig. 5).

There were only two gene regions that were mutated in lines started from multiple founders but not in lines started from the ancestor: *ompF/asnS* (in G3,5,6) and *rpsA* (in G3,5,6). In both cases mutations in these genes were also seen in at least once clone started from founder G1, which retained the ancestral *iclR* allele, indicating that these mutations are likely to confer some benefit in addition to that derived from any possible interaction with *iclR.* Similarly, there were two genes that were mutated in lines started from at least two different *iclR^Ev^* founders but not in any lines started founders that retained the ancestral *iclR* allele; ECB_01998 (G3 and 6 only) and *fadL* (G2 and 3 only). Mutations in these genes occurred in only a small number of lines, so cannot explain the general increase in evolvability or of *iclR* compensation.

To examine whether there were any general trends in mutated genes occurring in lines started from the same and different founders we examined two mutation similarity metrics. Intra-founder mutational parallelism was estimated as the number of observed mutated genes shared by clones started from each founder. Inter-founder convergence was estimated as the mean number of observed mutated genes shared among lines started from different founders. The log ratio of parallelism to convergence (PI:CI) therefore indicates the influence of the initial founder genotype in determining the identity of mutations selected during lactose adaptation.

We found that the amount of intra-founder parallelism varied, with some founders sharing many more mutated genes among derived clones than others (Fig 5). The PI:CI metric allowed us to test for an overall signal of mutational similarity depending on whether the founder had substituted an *iclR* mutation. When clones from the four *iclR^Ev^*founders were compared, the mean number of mutations in common was 1.11 (1.01-1.20 95%CI), indicating that, on average, each unique pairing of clones shared at least one mutated gene. However, the mean number of mutations in common among clones from *iclR^Anc^* founders and among all sequenced clones was similar at 0.97 (0.83-1.11 95%CI) and 1.07 (1.02-1.13 95%CI), respectively. Together these results indicate there is no signal of genetic parallelism dependent on the evolved *iclR* alleles. Instead, clones with and without evolved *iclR* alleles appear to be accessing similar mutational pathways in their adaptation to lactose.

## Discussion

Selection and analysis of compensatory mutations have a long history of use to determine the structure and function of molecular machines and regulatory pathways by revealing genes that interact with focal deleterious mutations (44). Compensatory mutations can also promote evolvability by influencing the form of the adaptive landscape (31, 45–47). We demonstrate a novel instance of this general phenomenon that depends on compensatory interactions that reverse the effect of conditionally deleterious mutations revealed upon an environmental shift. This reversal contributed to evolvability in lactose of populations started with at least four of six founder strains that had previously been selected in glucose.

At least three population genetic mechanisms could underlie the contribution of a ‘compensation pathway’ to increased evolvability of our founder strains. First, compensation could result in an increase in the beneficial mutation rate. This mechanism seems unlikely to explain our results. Indeed, even a ∼100-fold increase in the overall mutation rate conferred no significant benefit to our ancestral strain during adaption to a glucose limited environment (22, 48). Moreover, at least in the early period of adaptation to a new environment, the beneficial mutation rate does not appear to be limiting (7). Second, the effect size of available beneficial mutations could be greater in strains in which the cost of an *iclR* mutation has been compensated (46). However, the absence of any specific mutational signature is inconsistent with new large effect beneficial mutations becoming available.

In our view the most likely explanation for greater evolvability of the *iclR^Ev^* founders is simply that the *iclR*^Ev^ alleles have large deleterious effects in the lactose environment so that mutations that compensate that cost provide a single large fitness benefit relative to most beneficial mutations available to the ancestor. Two lines of evidence support this possibility. The mean effect of reverting evolved *iclR* mutations in founder strains was a 16.1% fitness benefit (±4.9% 95CI) (Table 1). To our knowledge, this represents a larger benefit than any other single mutation effect determined in any strain derived from our ancestor except for mutations in a key metabolic gene that had become depended on by subsequent adaptive changes (43, 49, 50). Additionally, the founder strains had mutations in several genes or gene regions—*nadR*, *rbs*, and *spoT*—that were also mutated in lines selected only in lactose and are likely to be associated with a fitness benefit (Table 1). Indeed, derivatives of the *iclR^Ev^*founders in which the *iclR* locus was reverted to the ancestral allele all had higher fitness than the ancestor. In other words, omitting *iclR* mutations, the net effect of mutations substituted in glucose was to confer a fitness increase in lactose. Compensating the cost of an *iclR* mutation would reveal this benefit.

In *E. coli*, IclR regulates transcription of the *aceBAK* operon (51). This operon encodes enzymes involved in the glyoxylate cycle that allows cells to directly split isocitrate to succinate and glyoxylate without the loss of CO_2_ (Lorca et al., 2007; Molina-Henares et al., 2006; Yamamoto and Ishihama, 2003). Knockout of IclR has been shown to increase expression of the *aceBAK* operon and increase cell density in some environments (52, 53). Notably, we found that *iclR^Ev^* alleles added directly to the ancestral REL606 strain did not have a significant impact on fitness in our glucose or lactose environments, indicating that epistatic interactions with other founder strain mutations influence the fitness effects of *iclR* alleles in both environments (Table 1). Unfortunately, except for mutations in *rbs,* which likely confer a benefit through preventing the expression cost of unnecessary gene products, we do not know the basis of the benefit conferred by mutations present in the founder strains (54). We therefore cannot predict how these mutations might interact to alter the effect of the *iclR^Ev^* mutations.

Whatever the mechanism through which compensation of *iclR*^Ev^ alleles confers increased evolvability, it is interesting to speculate whether a similar effect might generally affect genotypes with a history of fluctuating selection. Trade-offs, whereby adaptation to one environment is associated with fitness declines in another, are common though not ubiquitous (55, 56). For example, one comprehensive study found that adaptation of yeast to an environment buffered to pH 7.3 was associated with mean fitness declines in all of seven alternative environments, whereas adaptation to 21°C was associated with mean fitness increases to the same alternative environments (55). Notably, much work on trade-offs can determine only the overall effect of multiple substituted mutations. Even when the net effect of selection in one environment is a fitness increase in a second environment, some of the individual substituted mutations might be deleterious and therefore provide an opportunity to increase fitness through additional mutations that suppress their effect. However, simply compensating the effect of a mutation that becomes deleterious following an environmental shift would provide no direct advantage relative to genotypes that had not substituted the mutation. Rather, a benefit to compensation requires also that generally beneficial mutations are present and are being effectively masked by a conditionally deleterious mutation. That is, it is the distribution of correlated responses considered over all accumulated mutations that is important in predicting the potential for compensatory mutations to confer a benefit, not just the net correlated response. Determining the likelihood that a mix of generally beneficial and conditionally deleterious mutations are present in a given genotype requires detailed knowledge of the pleiotropic effects of individual substituted mutations that is not readily available.

Our results provide insight into a proposed mechanism of evolvability through compensation by confirming that mutations in *iclR* found among high evolvability founder strains were initially costly but evolved to become neutral or beneficial during continued selection of the founders in a new environment. The benefit of this change relative to naïve strains depends on the individual effects of previously selected mutations. Determining whether a history of fluctuating environmental selection can be expected to typically lead to increased evolvability requires an understanding of these individual effects.

## Acknowledgements

This work was supported by grant DEB-1253650 awarded to T.F.C. by the National Science Foundation, and by grant 19-MAU-082 awarded to T.F.C. by the Royal Society of New Zealand Marsden fund.

## Notes

### Competing Interest Statement

The authors have declared no competing interest.

## References

1. Payne JL, Wagner A. 2018. The causes of evolvability and their evolution. Nat Rev Genet 20:24–38.

2. Woods RJ, Barrick JE, Cooper TF, Shrestha U, Kauth MR, Lenski RE. 2011. Second-order selection for evolvability in a large *Escherichia coli* population. Science 331:1433–1436.

3. Phillips KN, Castillo G, Wünsche A, Cooper TF. 2016. Adaptation of *Escherichia coli* to glucose promotes evolvability in lactose. Evolution 70:465–470.

4. Gifford DR, Toll-Riera M, MacLean RC. 2016. Epistatic interactions between ancestral genotype and beneficial mutations shape evolvability in *Pseudomonas aeruginosa*. Evolution 70:1659–1666.

5. Hayden EJ, Ferrada E, Wagner A. 2011. Cryptic genetic variation promotes rapid evolutionary adaptation in an RNA enzyme. Nature 474:92–95.

6. Buckling A, Wills MA, Colegrave N. 2003. Adaptation limits diversification of experimental bacterial populations. Science 302:2107–2109.

7. Wünsche A, Dinh DM, Satterwhite RS, Arenas CD, Stoebel DM, Cooper TF. 2017. Diminishing-returns epistasis decreases adaptability along an evolutionary trajectory. Nat Ecol Evol 1:s41559-016–0061.

8. Barrick JE, Kauth MR, Strelioff CC, Lenski RE. 2010. *Escherichia coli rpoB* mutants have increased evolvability in proportion to their fitness defects. Mol Biol Evol 27:1338–1347.

9. Stiffler MA, Hekstra DR, Ranganathan R. 2015. Evolvability as a function of purifying selection in TEM-1 β-Lactamase. Cell 160:882–892.

10. Burch CL, Chao L. 2000. Evolvability of an RNA virus is determined by its mutational neighbourhood. Nature 406:625–628.

11. Couce A, Limdi A, Magnan M, Owen SV, Herren CM, Lenski RE, Tenaillon O, Baym M. 2024. Changing fitness effects of mutations through long-term bacterial evolution. Science 383:eadd1417.

12. Visser JAGMD, Cooper TF, Elena SF. 2011. The causes of epistasis. Proc Royal Soc B 278:3617–3624.

13. Arenas CD, Cooper TF. 2012. Mechanisms and selection of evolvability: experimental evidence. Fems Microbiol Rev 37:572–582.

14. Draghi JA, Parsons TL, Wagner GP, Plotkin JB. 2010. Mutational robustness can facilitate adaptation. Nature 463:353–355.

15. Fares M, Ruiz-González M, Moya A, Elena S, Barrio E. 2002. GroEL buffers against deleterious mutations. Nature 417:398.

16. Rajon E, Masel J. 2013. Compensatory evolution and the origins of innovations. Genetics 193:1209–1220.

17. Blount ZD, Barrick JE, Davidson CJ, Lenski RE. 2012. Genomic analysis of a key innovation in an experimental *Escherichia coli* population. Nature 489:513–518.

18. Quandt EM, Gollihar J, Blount ZD, Ellington AD, Georgiou G, Barrick JE. 2015. Fine-tuning citrate synthase flux potentiates and refines metabolic innovation in the Lenski evolution experiment. Elife 4:e09696.

19. Quandt EM, Deatherage DE, Ellington AD, Georgiou G, Barrick JE. 2014. Recursive genomewide recombination and sequencing reveals a key refinement step in the evolution of a metabolic innovation in *Escherichia coli*. Proc National Acad Sci 111:2217–2222.

20. Tanaka MM, Bergstrom CT, Levin BR. 2003. The evolution of mutator genes in bacterial populations: the roles of environmental change and timing. Genetics 164:843–854.

21. Rajon E, Masel J. 2011. Evolution of molecular error rates and the consequences for evolvability. Proc Natl Acad Sci USA 108:1082–1087.

22. Visser J de, Zeyl C, Gerrish P, Blanchard J, Lenski R. 1999. Diminishing returns from mutation supply rate in asexual populations 283:404–406.

23. Lynch M, Ackerman MS, Gout J-F, Long H, Sung W, Thomas WK, Foster PL. 2016. Genetic drift, selection and the evolution of the mutation rate. Nat Rev Genet 17:704–714.

24. Gerrish P, Colato A, Perelson A, Sniegowski P. 2007. Complete genetic linkage can subvert natural selection. Proc Natl Acad Sci USA104:6266–6271.

25. True H, Lindquist S. 2000. A yeast prion provides a mechanism for genetic variation and phenotypic diversity. Nature 407:477–483.

26. Tyedmers J, Madariaga ML, Lindquist S. 2008. Prion switching in response to environmental stress. Plos Biol 6:e294.

27. Rutherford SL, Lindquist S. 1998. Hsp90 as a capacitor for morphological evolution. Nature 396:336–342.

28. Zabinsky RA, Mason GA, Queitsch C, Jarosz DF. 2019. It’s not magic – Hsp90 and its effects on genetic and epigenetic variation. Semin Cell Dev Biol 88:21–35.

29. Bakerlee CW, Phillips AM, Ba ANN, Desai MM. 2021. Dynamics and variability in the pleiotropic effects of adaptation in laboratory budding yeast populations. Elife 10:e70918.

30. Remold S. 2012. Understanding specialism when the Jack of all trades can be the master of all. Proc Royal Soc B 279:4861–4869.

31. Szamecz B, Boross G, Kalapis D, Kovács K, Fekete G, Farkas Z, Lázár V, Hrtyan M, Kemmeren P, Koerkamp MJAG, Rutkai E, Holstege FCP, Papp B, Pál C. 2014. The genomic landscape of compensatory evolution. Plos Biol 12:e1001935.

32. Cooper TF, Lenski RE. 2010. Experimental evolution with *E. coli* in diverse resource environments. I. Fluctuating environments promote divergence of replicate populations. BMC Evol Biol 10:11.

33. Karkare K, Lai H-Y, Azevedo RBR, Cooper TF. 2021. Historical contingency causes divergence in adaptive expression of the *lac* operon. Mol Biol Evol 38:2869–2879.

34. Philippe N, Alcaraz J-P, Coursange E, Geiselmann J, Schneider D. 2004. Improvement of pCVD442, a suicide plasmid for gene allele exchange in bacteria. Plasmid 51:246–255.

35. Ferrieres L, Hemery G, Nham T, Guerout AM, Mazel D, Beloin C, Ghigo JM. 2010. Silent mischief: Bacteriophage Mu insertions contaminate products of *Escherichia coli* random mutagenesis performed using suicidal transposon delivery plasmids mobilized by broad-host-range RP4 conjugative machinery. J Bacteriol 192:6418–6427.

36. Phillips KN, Cooper TF. 2021. The cost of evolved constitutive *lac* gene expression is usually, but not always, maintained during evolution of generalist populations. Ecol Evol 11:12497–12507.

37. Lenski RE, Rose MR, Simpson SC, Tadler SC. 1991. Long-term experimental evolution in *Escherichia coli*. I. Adaptation and divergence during 2,000 generations. American Naturalist 138:1315–1341.

38. Deatherage DE, Barrick JE. 2014. Identification of mutations in laboratory-evolved microbes from next-generation sequencing data using breseq. Methods Mol Biology 1151:165–188.

39. Kryazhimskiy S, Rice DP, Jerison ER, Desai MM. 2014. Global epistasis makes adaptation predictable despite sequence-level stochasticity. Science 344:1519–1522.

40. Khan AI, Dinh DM, Schneider D, Lenski RE, Cooper TF. 2011. Negative epistasis between beneficial mutations in an evolving bacterial population. Science 332:1193–1196.

41. R Core Team. 2021. R: A Language and Environment for Statistical Computing.

42. Li G-M. 2008. Mechanisms and functions of DNA mismatch repair. Cell Res 18:85–98.

43. Quan S, Ray J, Kwota Z, Duong T, Balazsi G. 2012. Adaptive evolution of the lactose utilization network in experimentally evolved populations of *Escherichia coli*. PLoS Genetics 8:e1002444.

44. Bautista DE, Carr JF, Mitchell AM. 2021. Suppressor mutants: History and today’s applications. EcoSal Plus 9:eESP-0037-2020.

45. Covert AW, Lenski RE, Wilke CO, Ofria C. 2013. Experiments on the role of deleterious mutations as stepping stones in adaptive evolution. Proc Natl Acad Sci USA 110:E3171 8.

46. Helsen J, Voordeckers K, Vanderwaeren L, Santermans T, Tsontaki M, Verstrepen KJ, Jelier R. 2020. Gene loss predictably drives evolutionary adaptation. Mol Biol Evol 37:2989–3002.

47. Belinky F, Sela I, Rogozin IB, Koonin EV. 2019. Crossing fitness valleys via double substitutions within codons. BMC Biol 17:105.

48. Visser JAGMD, Rozen DE. 2005. Limits to adaptation in asexual populations. J Evol Biol 18:779–788.

49. Barrick JE, Yu DS, Yoon SH, Jeong H, Oh TK, Schneider D, Lenski RE, Kim JF. 2009. Genome evolution and adaptation in a long-term experiment with *Escherichia coli*. Nature 461:1243–1247.

50. Peng F, Widmann S, Wünsche A, Duan K, Donovan KA, Dobson RCJ, Lenski RE, Cooper TF. 2018. Effects of beneficial mutations in *pykF* gene vary over time and across replicate populations in a long-term experiment with bacteria. Mol Biol Evol 35:202–210.

51. Cortay JC, Nègre D, Galinier A, Duclos B, Perrière G, Cozzone AJ. 1991. Regulation of the acetate operon in *Escherichia coli*: purification and functional characterization of the IclR repressor. EMBO J 10:675–679.

52. Liu M, Ding Y, Chen H, Zhao Z, Liu H, Xian M, Zhao G. 2017. Improving the production of acetyl-CoA-derived chemicals in Escherichia coli BL21(DE3) through *iclR* and *arcA* deletion. BMC Microbiol 17:10.

53. Waegeman H, Beauprez J, Moens H, Maertens J, Mey MD, Foulquié-Moreno MR, Heijnen JJ, Charlier D, Soetaert W. 2011. Effect of *iclR* and *arcA* knockouts on biomass formation and metabolic fluxes in *Escherichia coli* K12 and its implications on understanding the metabolism of Escherichia coli BL21 (DE3). BMC Microbiol 11:70.

54. Cooper VS, Schneider D, Blot M, Lenski RE. 2001. Mechanisms causing rapid and parallel losses of ribose catabolism in evolving populations of *Escherichia coli* B. J Bacteriol 183:2834–2841.

55. Jerison ER, Ba ANN, Desai MM, Kryazhimskiy S. 2020. Chance and necessity in the pleiotropic consequences of adaptation for budding yeast. Nat Ecol Evol 4:601–611.

56. Chen V, Johnson MS, Hérissant L, Humphrey PT, Yuan DC, Li Y, Agarwala A, Hoelscher SB, Petrov DA, Desai MM, Sherlock G. 2023. Evolution of haploid and diploid populations reveals common, strong, and variable pleiotropic effects in non-home environments. eLife 12:e92899.

