## Supplemental Information for "Mutations in *iclR* increase evolvability by facilitating compensation that exposes cryptic beneficial mutations in experimental populations of *Escherichia coli*"

Running title: Exposing evolvability

Keywords: Evolvability, *iclR*, experimental evolution, adaptation, gene interactions

**Table S1** Bacterial strains and isolates used for strain construction.

| Strain ID | Alternate ID | Gen. Lactose Evolution | *iclR* allele | Reference |
| --- | --- | --- | --- | --- |
| Ancestor | REL606 | 0 | Ancestor | Lenski et al., 1991 |
| G2 | CD2 | 0 | T60P | Phillips et al. 2016 |
| G2-L1 | RKS88 | 1,000 | T60P |  |
| G3 | CD3 | 0 | G219R |  |
| G3-L1 | RKS94 | 1,000 | G219R |  |
| G5 | CD5 | 0 | -58/-142 |  |
| G5-L1 | RKS100 | 1,000 | -58/-142 |  |
| G6 | CD6 | 0 | Δ48 |  |
| G6-L1 | RKS108 | 1,000 | Δ48 |  |

**Table S2** Bacterial strains constructed during this study.

| Strain | Genetic background | *iclR* allele |
| --- | --- | --- |
| Anc-*iclR*_G2_ | Ancestor | T60P |
| Anc-*iclR*_G3_ | Ancestor | G219R |
| Anc-*iclR*_G5_ | Ancestor | -58 C→T |
| Anc-*iclR*_G6_ | Ancestor | Δ48 |
| G2-*iclR*_anc_ | G2 | Ancestor |
| G2-L1-*iclR*_anc_ | G2-L1 | Ancestor |
| G3-*iclR*_anc_ | G3 | Ancestor |
| G3-L1-*iclR*_anc_ | G3-L1 | Ancestor |
| G5-*iclR*_anc_ | G5 | Ancestor |
| G5-L1-*iclR*_anc_ | G5-L1 | Ancestor |
| G6-*iclR*_anc_ | G6 | Ancestor |
| G6-L1-*iclR*_anc_ | G6-L1 | Ancestor |

**Table** **S3** Primers used in this study.

| Primer  pair | | Sequence | Target size (bp) | Rest. Site* | Target *iclR* allele |
| --- | --- | --- | --- | --- | --- |
| IclR1 | F | 5’- AGAGTTCTGCAGCCGTAAAAGTTTCGGTGGAA -3’ | 833 bp | PstI | G219R and Δ48 |
|  | R | 5’- AGAGTTGAGCTCACAATGCAACAGCAGGGTTT -3’ |  | SacI |  |
| IclR2 | F | 5’- AGAGTTCTGCAGCGCTCAGTTGGGCTAAAAAG -3’ | 730 bp | PstI | T60P |
|  | R | 5’- AGAGTTGAGCTCTGTTCCACTTTGCTGCTCAC -3’ |  | SacI |  |
| IclR3 | F | 5’- AGAGTTCTGCAGCGCAGGATAGGGTGAACAAT -3’ | 792 bp | PstI | -58/-142 |
|  | R | 5’- AGAGTTGAGCTCGGTCGTGGAGTTGAAGGTGT -3’ |  | SacI |  |

*Restriction sites as noted were included in primers to facilitate directional cloning into the pDS132 vector.

**Table S4** Fitness effect of *iclR* alleles in founder and lactose evolved clones.

| Line | Evolved *iclR* allele | Gen. in lactose | Fitness with evolved *iclR*  allele | Fitness with ancestral *iclR*  allele | *df* | *t*-value | P-value* |
| --- | --- | --- | --- | --- | --- | --- | --- |
| G2 | T60P | 0 | 0.97 | 1.14 | 21.6 | 3.58 | 0.001 |
|  |  | 1,000 | 1.42 | 1.42 | 11.1 | 0.04 | 0.486 |
| G3 | G219R | 0 | 1.03 | 1.22 | 12.9 | 9.31 | <0.001 |
|  |  | 1,000 | 1.37 | 1.36 | 17.9 | 0.24 | 0.405 |
| G5 | -58 C→T | 0 | 1.04 | 1.12 | 12.4 | 1.66 | 0.061 |
|  |  | 1,000 | 1.36 | 1.46 | 8.2 | 1.52 | 0.083 |
| G6 | Δ46 | 0 | 1.03 | 1.24 | 10.9 | 3.80 | 0.001 |
|  |  | 1,000 | 1.32 | 1.32 | 5.8 | 0.12 | 0.454 |

*^*^*One-tailed P-values testing the prediction that evolved *iclR* alleles are costly in the lactose environment.
